# PRC2-mediated H3K27me3 uncouples PHE1 binding from transcriptional activation at imprinted genes

**DOI:** 10.64898/2026.11.19.767895

**Authors:** Heinrich Bente, Melanie Höhne, Claudia Köhler

**Affiliations:** Department of Plant Reproductive Biology and Epigenetics, Max Planck Institute of Molecular Plant Physiology, Potsdam 14476, Germany

## Abstract

Genomic imprinting results in parent-of-origin-dependent gene expression and plays an important role in endosperm development. The type I MADS-box transcription factor PHERES1 (PHE1) has been implicated in the regulation of imprinted genes, particularly paternally expressed genes (PEGs). Silencing of the maternal alleles of PEGs is mediated by repressive histone modifications, including H3K27me3 established by the Polycomb Repressive Complex 2 (PRC2). Here, we established an adapted low-input CUT&Tag protocol to profile the allele-specific genome-wide binding of PHE1 in the endosperm and compared the resulting binding profiles with previously published ChIP-seq data. Our CUT&Tag analysis substantially recapitulated the previously identified PHE1-binding landscape while requiring considerably less input material. Consistent with earlier findings, PHE1 binding was enriched at RC/Helitron transposable elements and imprinted genes, specifically PEGs. Interestingly, most PHE1-bound PEGs exhibited binding on both maternal and paternal alleles despite H3K27me3 enrichment on their maternal alleles. Perturbation of H3K27me3 deposition did not alter the predominantly biallelic PHE1 occupancy observed at these loci, supporting that PHE1 binding can persist in an H3K27me3-associated chromatin environment. In contrast, analysis of gene expression in *fertilization independent seed* (*fis*) *prc2* mutants impaired in H3K27me3 establishment caused increased expression of PHE1 targets and altered their parent-of-origin-specific expression patterns. Our findings reveal that biallelic transcription factor occupancy can coexist with parent-of-origin-specific transcriptional regulation, highlighting the importance of chromatin context in determining the transcriptional output of PHE1-bound genes.

## Introduction

The endosperm is a nourishing tissue that supports embryo growth. It is formed after double fertilization, whereby one sperm cell fuses with the central cell to form the endosperm, while the other fertilizes the egg to develop into the embryo. In *Arabidopsis thaliana* (hereafter referred to as Arabidopsis), fertilization of the homodiploid central cell by a haploid sperm cell gives rise to the triploid endosperm, characterized by a 2:1 maternal-to-paternal (2m:1p) genome ratio. Maintaining this ratio is essential for normal endosperm development, as deviations, such as those resulting from crosses between parents of different ploidy levels, frequently leads to seed abortion (Dilkes et al., 2008, Scott et al., 1998).

The endosperm is also a major site of genomic imprinting, an epigenetic phenomenon that results in parent-of-origin-specific gene expression (Gehring et al., 2011, Hsieh et al., 2011, Batista and Köhler, 2020). Depending on the parental origin of the expressed allele, imprinted genes are classified as maternally expressed genes (MEGs) or paternally expressed genes (PEGs). The non-expressed allele is typically silenced by repressive epigenetic modifications, collectively referred to as the ‘imprint’. The imprint varies between the affected parental allele, whereby MEGs are typically characterized by reduced DNA methylation on the maternal allele, as a result of DNA demethylation of small repetitive regions in the central cell (Gehring et al., 2009, Gehring et al., 2011, Pignatta et al., 2014, Hsieh et al., 2011). On the contrary, the majority of PEGs are marked by trimethylation at lysine 27 of histone H3 (H3K27me3) on their maternal allele, mediated by the activity of the FERTILIZATION INDEPENDENT SEED (FIS) Polycomb Repressive Complex 2 (PRC2) in the central cell (Moreno-Romero et al., 2016, Mozgova et al., 2015, Zhang et al., 2014).

The M-type MADS-box transcription factor PHERES1 (PHE1) has previously been shown to bind to numerous key regulators of endosperm development, including many PEGs) (Batista et al., 2019. Although loss of PHE1 and its paralog PHE2 does not cause detectable phenotypic abnormalities in diploid seeds, the *phe1 phe2* double mutant suppresses triploid seed abortion (Batista et al., 2019), supporting a causal role for increased PEG expression in triploid seed abortion and a role of PHE1 in activating of these genes (Kradolfer et al., 2013, Wolff et al., 2015, Huang et al, 2017). Previous work proposed that PHE1 binds to both parental alleles of PEGs, indicating that allele-specific H3K27me3 prevents activation of the maternal alleles despite PHE1 presence. However, this conclusion was based on the analysis of two PEGs, limiting its general applicability. A genome-wide investigation of allele-specific PHE1 binding was previously hampered by the large amounts of developmental stage-specific hybrid material required for ChIP-seq, the gold-standard method for mapping transcription factor binding sites at the time. To overcome this limitation, we adapted the Cleavage under target and tagmentation (CUT&Tag) protocol (Kaya-Okur et al., 2019), which requires substantially less input material than conventional Chip-seq. Using samples comparable to those analyzed in previous Chip-seq studies, we demonstrate that our CUT&Tag protocol identified a largely overlapping set of PHE1 targets, including a high proportion of PEGs. Allele-specific analysis revealed that PHE1 binds to both parental alleles of the majority of its targets genome-wide, including PEGs. We further show that this biallelic binding occurs independently of H3K27me3 deposition, whereas H3K27me3 prevents transcriptional activation of the maternal alleles, despite PHE1 binding. Together, our findings provide new insights into the interplay between MADS-box transcription factors and epigenetic imprinting mechanisms in regulating seed development. Moreover, our adapted CUT&Tag protocol provides a powerful approach for profiling genome-wide transcription factor binding in tissues with severely limited sample availability, such as the endosperm.

## Results

### CUT&Tag works reliably for an endosperm-expressed transcription factor

The transcription factor PHE1 is transiently expressed in the endosperm for only approx. 5 days after pollination (DAP) (Weinhofer et al., 2010). This temporally and spatially restricted expression makes it difficult to generate sufficient amounts of time-resolved input material for traditional methods such as ChIP-seq for the identification of transcription factor binding sites. To overcome this challenge, we have adapted the recently published CUT&Tag protocol (Kaya-Okur et al., 2019), proposed to require only a fraction of previously needed input material. To evaluate the performance of CUT&Tag for PHE1-GFP compared to ChIP-seq, we generated CUT&Tag libraries from the same transgenic lines at comparable time points of two DAP, as had been used in previous work (Batista et al., 2019). A genome-wide analysis of our newly generated CUT&Tag data of *PHE1::PHE1-GFP* containing plants data revealed good Spearman correlation between the samples from independent biological and methodological replicates for both PHE1-GFP and anti-H3 sample (Supplementary Fig. S1A). To compare the CUT&Tag target enrichments in comparison to ChIP-seq, we first observed well-known PHE1 targets manually in a genome browser using three independent replicates, in comparison to anti-H3 samples (for CUT&Tag) or input samples (for ChIP-seq) as background control (Fig. 1A). The upstream regions of known PHE1 target genes such as *YUCCA10 (YUC10) AGAMOUS-LIKE 23 (AGL23)* and *HAIKU2 (IKU2)* showed stronger signal from anti-GFP CUT&Tag samples (dark green), but not in anti-H3 CUT&Tag (dark grey), revealing enrichment for well-known targets (Fig. 1A). Importantly, CUT&Tag performed on wild-type siliques containing no *PHE1::PHE1-GFP* transgene did not show enrichment of known PHE1 target regions (Fig. 1A). Compared to ChIP-seq data (Batista et al., 2019) the signal enrichment was detectable in the same promotor region, however, CUT&Tag led to a much more prominent enrichment (Fig. 1A). A stronger target-enrichment through CUT&Tag had previously also been reported for histone modifications in comparison to ChIP-seq (Kaya-Okur et al., 2019, Abbasova et al., 2025). To obtain only high confidence CUT&Tag peaks, we adopted the ENCODE project pipeline including the use of permissive peak calling followed by pairwise irreproducible discovery rate (IDR) peak ranking (Supplementary Fig. S1B), which resulted in 1090 peaks. Different replicates showed good to moderate Spearman correlation over these 1090 regions, ranging from 0.53 to 0.76 depending on replicate combination (Supplementary Fig. S1C). In contrast, transgene-free samples resulted in 10 enriched regions, of which only 8 overlapped with the 1090 PHE1 target regions (Supplementary Fig. S1D). The fraction of fragment in peaks (FFiPs) over these 1090 regions was approximately five-fold higher in anti-GFP libraries from *PHE1::PHE1-GFP* containing samples, compared to Col-0 derived anti-GFP samples (Supplementary Fig. S1E). Annotation of PHE1 targets from both methods led to unique target genes, in total 994 for CUT&Tag and 1749 for ChIP-seq (Fig. 1B), with a significant number of 562 PHE1 target genes (Hypergeometric test p < 2.2*10^-16^) were shared between CUT&Tag and ChIP-seq (Fig. 1C). The discrepancy in identified target-regions is a consequence of the higher number of CUT&Tag replicates compared to the previous ChIP-seq data, allowing the more stringent peak selection inspired by the ENCODE IDR workflow.

**Figure 1:**
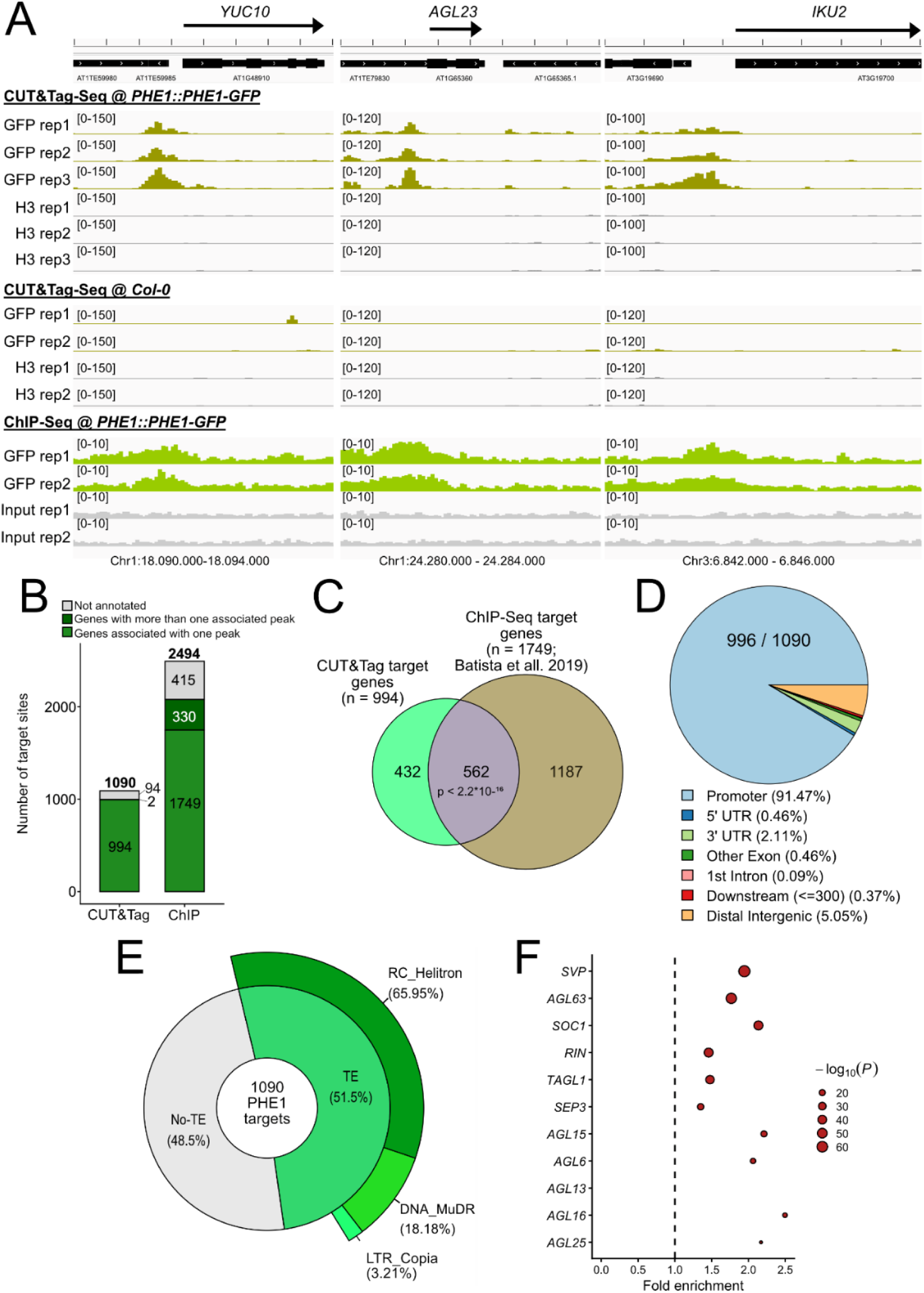
CUT&Tag reveals known PHE1 binding sites. A) Browsershots of replicates of known PHE1 targets as identified by Batista et al 2019. Tracks are shown from CUT&Tag-seq in Col-0 *PHE1::PHE1-GFP* containing plants and Col-0 transgene-free wild-type plants and ChIP-Seq tracks from Batista et al. 2019. Shown are three independent replicates obtained from GFP (green) and background (grey, H3 = dark grey, ChIP input = light grey) antibody samples. Reads where normalized to reads per genomic content (RPGC), per track range indicated in brackets. B) Number of annotated peaks in CUT&Tag and ChIP-seq data. C) Overlap of uniquely annotated genes between CUT&Tag and ChIP-seq targets. Statistical significance was tested via hypergeometric test. D) Piechart of CUT&Tag peak feature annotations per ChIPseeker using a -1500 bp + 500 bp promoter relative to TSS. E) Piedonut chart depicting the relative overlap of PHE1 CUT&Tag targets and transposable elements (TEs). F) Enrichment of MADS motives in uniquely annotated CUT&Tag peaks as determined by Homers findMotifsGenome.pl.

We performed feature annotation of the CUT&Tag-identified 1090 PHE1 target sides, which showed that 91.47% of PHE1 target sites are located in promoter regions (Fig. 1D, -1.5 to +0.5 kb relative to TSS), as previously shown for transcription factors (Yu et al., 2016, Heyndrickx et al., 2014, O’Malley et al., 2016). Similar annotation of the ten target regions from wild-type samples, showed that only five of those falling into the same promoter window (Supplementary Fig. S1E). Enriched regions identified by CUT&Tag displayed similar features as those identified by ChIP-seq experiments, including enrichment for transposable elements (TEs), specifically for RC/Helitrons (Batista et al., 2019). CUT&Tag confirmed that half (52%) of PHE1 target regions overlap with TEs, of which 66% belong to the RC/Helitron superfamily (Fig. 1E). Thus, 34% of all PHE1 target sides overlap with RC/Helitrons, corresponding to previous ChIP-seq experiments (Batista et al., 2019). Lastly, as PHE1 is a MADS box transcription factor, we tested if MADS motives were prevalent in the 994 target sides. We found enrichment for multiple MADS motives, among them the previously identified SVP MADS motif (Fig.1 F). Overall, the in-depth comparison between CUT&Tag and ChIP-seq performed in comparable tissues, revealed that both methods could recover promoter-target regions with good reproducibility, while CUT&Tag library preparation only required a fraction of the input. Nevertheless, less peaks were recovered with CUT&Tag after stringent analysis, but these peaks were more prominent compared to peaks identified by ChIP-seq.

### Imprinted regulation of PHE1 targets is a consequence of allele-specific H3K27me3

In the previous ChIP-seq experiment, many PHE1 targets were imprinted genes (Batista et al., 2019). Among the PHE1 targets established using CUT&Tag, we also found a substantial number of imprinted genes, specifically PEGs. We opted to verify the enrichment of imprinted genes in our CUT&Tag data using a list of high confidence imprinted genes, appearing in at least two independent studies referred to as ‘core’ imprinted genes (Del Toro-De Leon et al., 2024). This list consists out of 268 ‘core’ MEGs as well as 103 ‘core’ PEGs, we simply refer to as MEGs or PEGs respectively. CUT&Tag revealed that only a few MEGs (6/268, 2.24%) were targeted by PHE1, whereas a significant number of 20 PEGs (hypergeometric test p = 2.6*10^-11^), thus 19.42% of PEGs were PHE1 targets (Fig. 2A). This high number of targeted PEGs raised the question if PHE1 targeting itself exerts allele specific binding. To investigate allele-specificity, we made use of single nucleotide polymorphism between the natural Arabidopsis accessions Col-0 and Landsberg *erecta* (L*er*). We used L*er* as maternal parents whereas *PHE1::PHE1-GFP* transgenic plants in Col-0 background were used as pollen donor. To allow comparability with publicly available imprinting data, F1 siliques were harvested at 4DAP and subjected to CUT&Tag sequencing. However, PHE1 PEG targets did not show allele-specific PHE1 binding across all three replicates (Fig. 2B), supporting our previous observations that were based on two analyzed PEGs (Batista et al., 2019). Similarly, the identified MEG targets showed mainly biallelic PHE1 occupancy (Fig. 2B), in contrast to previously detected predominantly maternal binding (Batista et al., 2019). However, the two previously analyzed MEGs were not detected in the CUT&Tag data, leaving the possibility that allele-specific binding of PHE1 may occur at some MEGs. Browershots of PHE1 PEG targets confirmed the biallelic signal in the PHE1 target region (Fig. 2C). Consistent with the trend for PEGs/MEGs, the majority of non-imprinted PHE1 targets were bound on the maternal and paternal alleles with an average maternal ratio of 0.72, close to the expected biallelic ratio of 0.66 (2m:1p) in the endosperm (Fig. 2D). Similarly, the maternal ratio in defined promoter windows (-1500 bp TSS +500 bp) of all PHE1 target genes showed a mean maternal ratio of 0.76 (Supplementary Fig. S2). The strong biallelic binding of PHE1 contrasts with the paternally biased expression of its target genes, raiding the question how maternal alleles remain silenced despite PHE1 binding.

**Figure 2:**
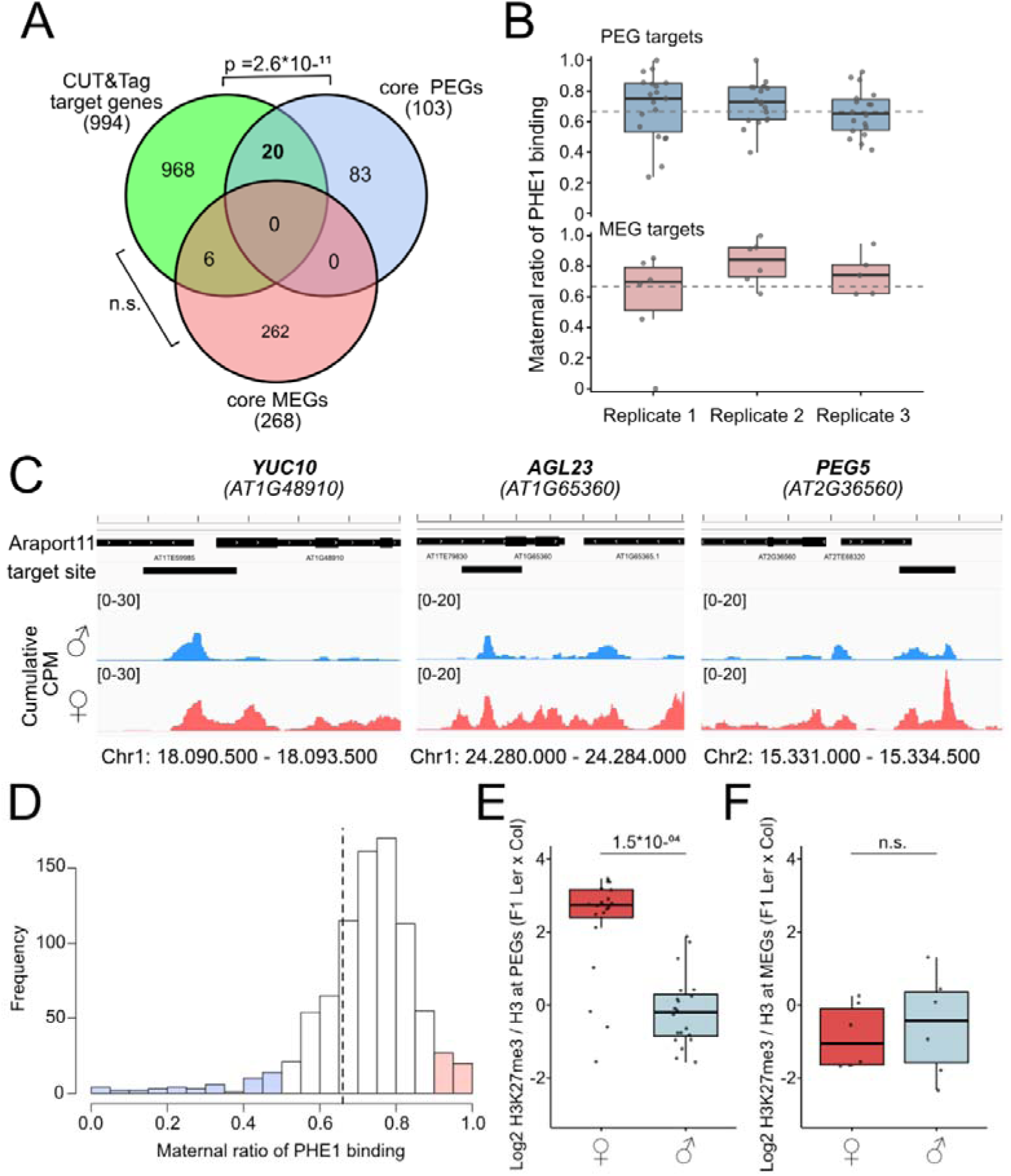
Allele-specific CUT&Tag reveals biallelic binding of PHE1 to the majority of its targets. A) Venn diagram showing the overlap of PHE1 targets, with "core PEGs" and "core MEGs" as defined in Del Toro de León et al. 2024. Statistically significant overlaps were determined via hypergeometric test. B) Maternal ratio of PHE1 binding on targeted core imprinted genes established via CUT&Tag on PHE1 target sides with sufficient numbers of informative reads (>5 reads/SNP). C) Browsershot of PHE1 PEG targets showing biallelic binding. Tracks were normalized to CPM normalized to the combined allele-specific library size (cumulative CPM, see material and methods). D) Histogram of maternal ratios from all PHE1 target sites with sufficient informative reads (>5 reads/SNP). Targets with allele-bias are marked by color (maternal ratio < 0.5, blue); for maternal allele (maternal ratio > 0.9, red); or bialleic PHE1 binding (maternal ratio >=0.5 & <= 0.9, white). E) Allele-specific log2 of H3K27me3 / H3 on PEGs targeted by PHE1 in endosperm nuclei 4 DAP in F1 Ler x Col. F) Allele-specific log2 of H3K27me3 / H3 on MEGs targeted by PHE1 in endosperm nuclei 4 DAP in F1 Ler x Col. Statistical difference was determined with two-sided Wilcoxon signed-rank test p < 0.05. Endosperm-specific H3K27me3 and H3 data are from Del Toro de León et al. 2024.

We previously found that PEGs are associated with maternal-specific H3K27me3 (Moreno-Romero et al., 2019, Moreno-Romero et al., 2016), suggesting that the parent-specific presence of H3K27me3 is decisive for allele-specific expression of PHE1 target genes rather than allele-specific binding of PHE1. We tested this hypothesis by analyzing the allele-specific presence of H3K27me3 at PHE1 PEG target genes and found indeed that the maternal alleles of PHE1 PEG targets are enriched for H3K27me3 (Fig. 2E), suggesting that the presence of H3K27me3 determines whether PHE1 binding leads to expression of the targeted allele. We did not detect allele-specific H3K27me3 for the 6 MEG PHE1 targets, consistent with previous work showing that imprinting of most MEGs is independent of allele-specific H3K27me3 (Fig. 2F) (Moreno-Romero et al., 2019, Moreno-Romero et al., 2016). Also, consistent with previous observations, PHE1-binding sites on paternal MEG alleles exhibited increased CG DNA methylation; however, due to the limited number of PHE1-targeted MEGs, this difference was not statistically significant (Supplementary Fig. S3A). In contrast, PEGs targeted by PHE1 exhibited increased CG methylation on paternal alleles and, consistent with previous findings, increased CHG methylation on maternal alleles (Supplementary Fig. S3B) (Moreno-Romero et al., 2019). Together, these results suggest that PHE1 binds the majority of its targets biallelically, including imprinted genes. For PEGs, paternal-allele specific expression is therefore likely maintained through H3K27me3-mediated silencing of the maternal allele.

### PHE1 binding is compatible with allele-specific H3K27me3

Publicly available endosperm-specific expression data (Moreno-Romero et al., 2019) showed that the average PHE1 target expression is increased in comparison to all annotated genes in Araport11, and slightly increased compared to expressed genes in the endosperm (RPKM >5, Fig. 3A). Furthermore, PHE1 targets had slightly, but significantly higher levels of H3K27me3 compared to all endosperm expressed genes (Fig. 3B). This enrichment of H3K27me3 compared to H3 was not only prevalent in the gene body region, but also in the promoter region where PHE1 binds (Fig. 3C). To explore a possible effect of H3K27me3 on PHE1 binding, we used kmeans clustering to obtain three clusters around the midpoint of all 994 PHE1 targets in a +-1kb interval. This resulted in cluster 1 containing 189 PHE1 target sites enriched in H3K27me3, cluster 2 with 410 PHE1 target sides with lower levels of H3K27me3, and cluster 3 with 395 genes having the least H3K27me3 (Fig. 3D). Cluster 1 contained 16 of the 20 PHE1 PEG targets, whereas the remaining four PEGs belonged to cluster 2 (Fig. 3E), consistent with previous work revealing that PEGs are marked by H3K27me3 (Moreno-Romero et al., 2019, Moreno-Romero et al., 2016). Most H3K27me3 in cluster 1 is present on the maternal alleles (Fig. 3F), which was a general trend and not restricted to PEGs and applied for both directions of the crosses (Supplementary Fig. S4A). Similar as for PEGs (Supplementary Fig. S4B-C) the presence of H3K27me3 reduced expression of the maternal alleles, biasing genes located in cluster 1 significantly for paternal expression, whereas genes in clusters 2 and 3 were biallelically expressed (Fig. 3G). Nevertheless, genes in cluster 1 were biallelically bound by PHE1 (Fig. 3H), despite the presence of maternal H3K27me3 on PHE1 target sites.

**Figure 3:**
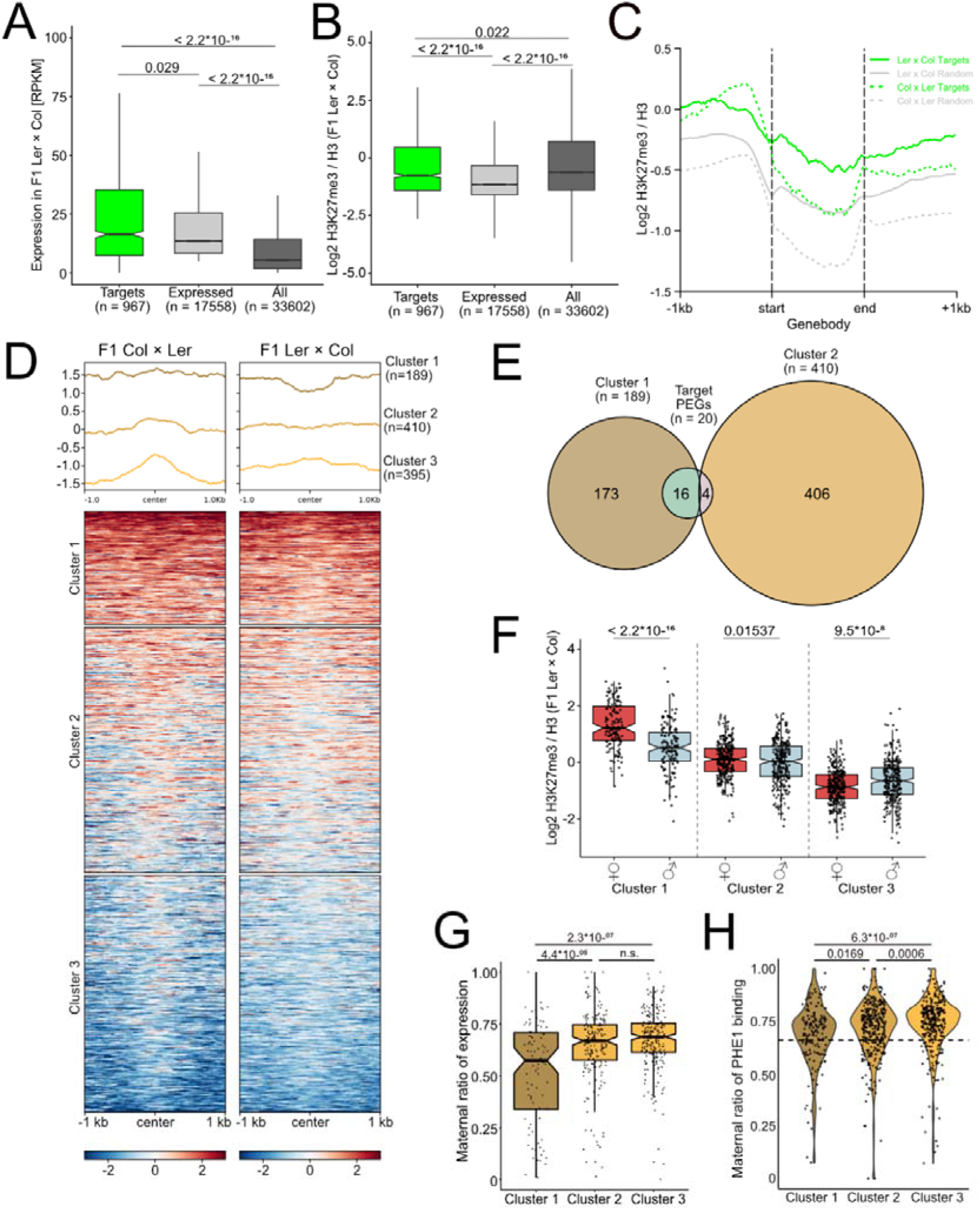
PHE1 targets are enriched for H3K27me3 on gene bodies and promoter regions on the maternal alleles. A) Expression of PHE1 targets in comparison to all expressed genes (RPKM >5) and all annotated genes. Expression data are from INTACT purified endosperm nuclei at 4 DAP in F1 Ler × Col seeds form Moreno Romero et al. 2019. B) Log2 normalized levels of total H3K27me3/H3 levels at PHE1 target genes, at all expressed genes in the endosperm in F1 Ler x Col crosses. C) Metagene plot over PHE1, targets and random genes in reciprocal crosses. D) Kmeans clustering and heatmap over all PHE1 target sides in a window of 1 kb up- and downstream to mid-point of annotated peaks. E) Venn diagram depicting the overlap of PHE1 PEG target genes with identified clusters. F) Parental-specific H3K27me3 in clusters as defined before. Statistical difference was determined with two-sided Wilcoxon signed-rank test p < 0.05.G) Maternal ratio of PHE1 target gene expression per cluster at 4 DAP. Expression data from Moreno-Romero et al. 2019. H) Maternal ratio of PHE1 binding by clusters. Wilcoxon rank-sum test was applied in panels A,B, G and H. H3K27me3 data are from Del Toro-De León et al. 2024.

To further test whether the presence of H3K27me3 on the maternal alleles does not interfere with PHE1 binding, we tested PHE1 binding in seeds of *mea/+* mutants, where we introduced the *PHE1::PHE1-GFP* through pollen. *Mea* is a maternal gametophytic mutant, with 50% of seeds from a *mea/+* plant being depleted for MEA function and abort (Grossniklaus et al., 1998). We compared the allele-specific PHE1 binding at known PHE1 target sites (Fig. 2). (Fig. 4A). Consistent with the data showing PHE1 binding in the presence of H3K27me3 (Fig 3D), PHE1-binding on the maternal alleles was not substantially increased in the *mea* mutant (Fig. 4A-C). While the ratio of bound alleles was significantly different for genes in cluster 1, there was no pronounced change towards PHE1 binding of the maternal allele (Fig. 4A). This suggests that while *mea/+* indeed influences PHE1 binding, this affects both alleles not just the maternal one. One possible explanation could be that increased expression of *PHE1* in the *mea* mutant (Köhler et al., 2003) leads to increased expression of PHE1 targets.

**Figure 4:**
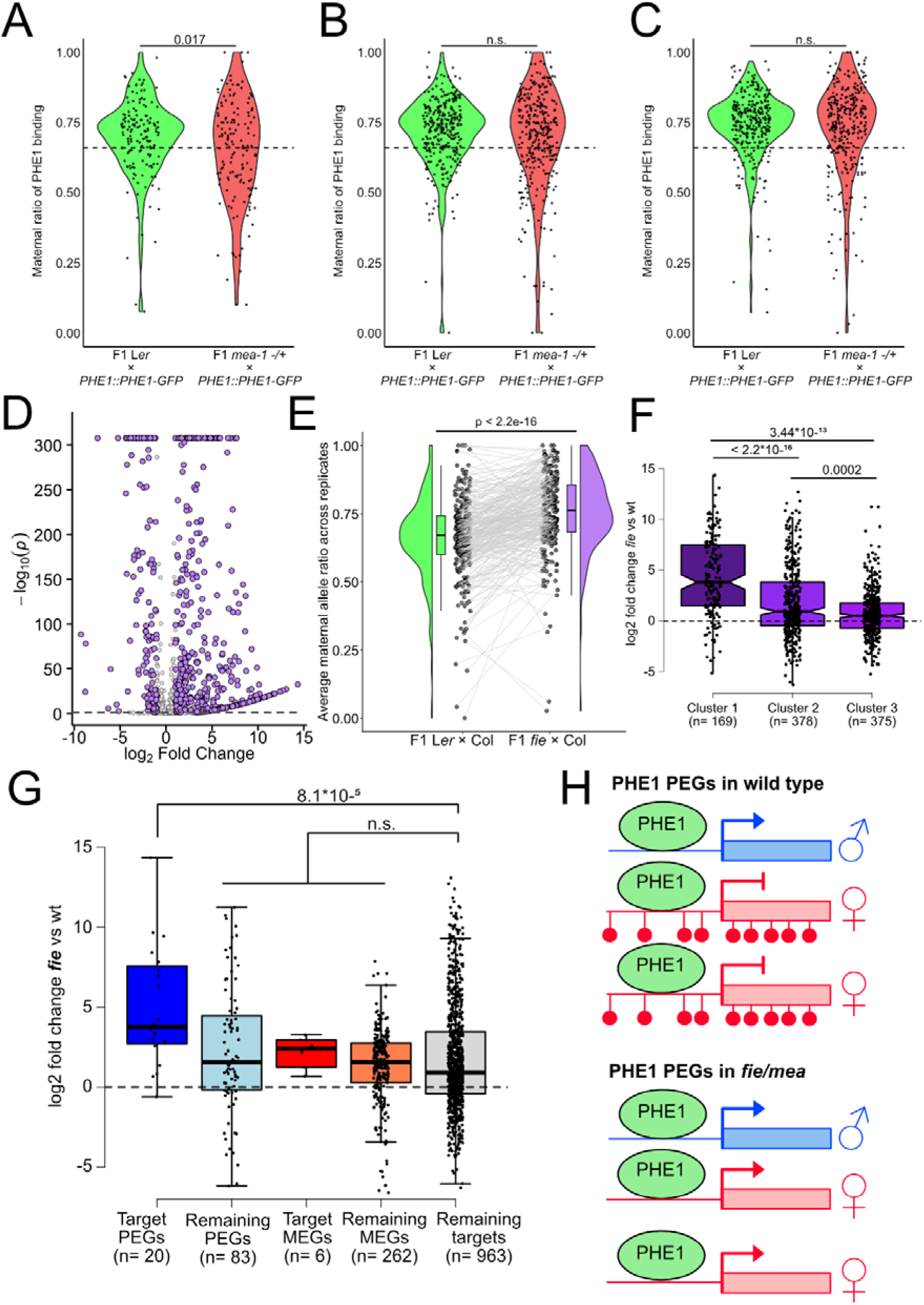
PHE1 targets are repressed by H3K27me3. A) Violin plots depicting the allele-specific PHE1 binding in Ler x *PHE1::PHE1-GFP* (Col) in afore established cluster 1 (A) cluster 2 (B) or cluster 3 (C) at PHE1 target sides. Dashed lines represent the theoretical parental ratio of 2m:1p, 0.66D) Volcano plot of PHE1 targets genes identified by CUT&Tag in FIE deficient seeds. E) Rainclould plot of allele ratios from PHE1 target genes meeting allele criteria (>5 reads per SNP, see material and methods) in F1 endosperm of Ler x Col and F1 *fie* x Col. Mean maternal allele ratios were compared between genotypes using a two-sided Wilcoxon signed-rank test. F) Differential gene regulation in H3K27me3 clusters as defined earlier.G) Log2-fold change between *fie* and wt crosses of PEGs, MEGs (PHE1 targets and all) as well as all target genes. Transcriptome data are from Hsieh et al. 2011. Two-sided Wilcoxon signed-rank tests were used for panels A,B,C,E and two-sided Wilcoxon rank-sum tests were applied in panels F and G; p < 0.05. H) Modell of PHE1 binding at PEGs in wild-type and H3K27me3 compromised conditions. Allele-specific expression is indicated by arrows, thicker arrows indicate stronger expression. Paternal alleles are marked in blue, maternal in red. Repressive H3K27me3 are illustrated by red upside-down lollipops.

To test this hypothesis, we analyzed publicly available endosperm transcriptome data from plants deficient in FIE, a core component of the FIS-PRC2, which also comprises MEA, FIS2 and MSI1 (Hsieh et al., 2011, Köhler et al., 2003). Many PHE1 target genes were upregulated in *fie* compared to wild type (Fig. 4D). The upregulation of PHE1 target genes in *fie* was accompanied by stronger expression of the maternal alleles, consistent with H3K27me3 being predominantly located on maternal alleles (Fig. 4E). This was also reflected by PHE1 target genes located in cluster 1 (having highest H3K27me3) showing the strongest expression changes in *fie*, followed by genes in cluster 2, whereas cluster 3 showed only minor upregulation compared to wild type (Fig. 4F). These data suggest that genes with higher H3K27me3 enrichment exhibit stronger transcriptional upregulation in FIS-PRC2 mutants, suggesting that the extent of H3K27me3 enrichment is associated with the magnitude of PRC2-dependent transcriptional repression. PHE1 targeted PEGs were among the most highly upregulated genes in *fie* and were significantly higher expressed compared to all other imprinted genes and also non-imprinted PHE1 target genes (Fig 4G). Notably, PHE1-targeted PEGs, which showed the strongest transcriptional upregulation, also exhibited the highest levels of H3K27me3 enrichment on their maternal alleles compared with the other gene categories. Interestingly, the strong upregulation of PHE1 targeted PEGs was not accompanied by increased PHE1 targeting to the maternal alleles (Fig. 4A-C). This suggests that PHE1 binds both parental alleles, but H3K27me3 prevents transcriptional activation of the maternal alleles, which become derepressed upon loss of FIS-PRC2 activity. Collectively, these results demonstrate that PHE1 binds to both alleles of several H3K27me3-marked genes. H3K27me3 depletion results in increased expression of PHE1 target genes, predominantly driven by enhanced maternal allele expression, consistent with the preferential deposition of H3K27me3 on maternal alleles in the endosperm (Moreno-Romero et al., 2016). Together, these findings suggest that H3K27me3 restricts PHE1-mediated transcriptional activation, particularly on maternal alleles, thereby contributing to the maintenance of parent-of-origin-specific gene expression.

## Discussion

Here we report CUT&Tag as a quick and low-cost alternative for genome-wide monitoring of TF binding sites from limited amount of tissue, like the Arabidopsis endosperm. Using this technique, we were able to monitor allele-specific binding of the PHE1, a key regulator of PEGs in Arabidopsis and to compare this data with our previously established ChIP-seq data of PHE1 from the same tissue (Batista et al., 2019). Compared to ChIP-seq, CUT&Tag comes with several advantages that are particularly relevant when working with limited amounts of tissue. CUT&Tag protocols are substantially faster than ChIP-seq, require less input material, and generally result in better signal to noise output, obtained from less deep sequenced samples. Due to the shorter and hence faster protocols, CUT&Tag-seq, as well CUT&RUN-seq are slowly replacing ChIP-seq, especially for the analysis of histone marks in mammalian cell systems (Kaya-Okur et al., 2019, Bartosovic et al., 2021), but also in plants (Zheng and Gehring, 2019, Tao et al., 2020, Del Toro-De Leon et al., 2024).

Still, CUT&Tag is rarely used to obtain genome-wide binding events of transcription factors, and it is often recommended to use CUT&RUN instead, as the general higher salt concentrations in CUT&Tag buffers, which are needed to dampen the DNA affinity of free pA-Tn5, are supposed to prevent weak TF-DNA interactions (Kaya-Okur et al., 2020). Indeed, our CUT&Tag against PHE1-GFP identified about half the number of the previously identified ChIP-seq targets (Batista et al., 2019); with about half of those identified targets being shared between both methods. Similarly, a recent comparison of CUT&Tag-seq- and ChIP-seq-mediated profiling of histone modifications found about half of the target sites overlapped between both methods (Abbasova et al., 2025), similar to the overlap found in this study. Together with the significant overlap between CUT&Tag and ChIP-seq targets, the enrichment of MADS-box binding motifs and TEs, particularly RC/Helitrons, supports the reliability of CUT&Tag in identifying PHE1 binding sites. Consistent with previous PHE1 ChIP-seq experiments (Batista et al., 2019), we found a significant enrichment of imprinted genes, particular PEGs, among PHE1 target genes identified by CUT&Tag. The lower input requirement of CUT&Tag over ChIP-seq enabled us to generate allele-specific PHE1 binding profiles, thus we were able to test if parent-of-origin-specific expression is accompanied by allele-specific PHE1 binding. We were able to show that all by CUT&Tag identified PEG PHE1 targets, PHE1 localizes to both alleles, the actively transcribed paternal, but also to the repressed maternal allele. These data support and extend our previous findings that were based on two PEG PHE1 targets (Batista et al., 2019). PEGs in the Arabidopsis endosperm are characterized by the presence of H3K27me3 on the maternal alleles (Moreno-Romero et al., 2019, Moreno-Romero et al., 2016). We found that PHE1 binds to the repressed maternal alleles of PEGs, but also of other, not imprinted genes marked by H3K27m3 on their maternal alleles and surrounding the PHE1 binding sites. Repressive H3K27me3 found at PEGs are established in the central cell by the FIS-PRC2 and inherited to the endosperm (Gehring, 2013, Rodrigues and Zilberman, 2015). Since *PHE1* is expressed after fertilization (Köhler et al., 2005), it is unlikely that PHE1 itself recruits the FIS-PRC2, while we cannot rule out a maintenance function. Possibly the function of PHE1 depends on different interaction partner, as MADS box TFs show a broad interaction matrix with each other (Bemer et al., 2010, de Folter et al., 2005).

Consistent with our finding that PHE1 shows binding to regions marked by H3K27me3, we found that depletion of this modification in *mea* mutants did not impact on PHE1 binding. However, the maternal alleles of PHE1 targets became activated upon H3K27me3 depletion, revealing that the presence of H3K27me3 prevents activation of the maternal alleles despite PHE1 binding. Interestingly we observed a positive connection between H3K27me3 enrichment at PHE1 targets and the magnitude of gene activation following PRC2 loss. Previous work established that the level of H3K27me3 required to maintain transcriptional silencing depends on promoter activity, with transcriptionally poised promoters becoming active despite retaining levels of H3K27me3 (Jadhav et al., 2020). What determines the level of H3K27me3 remains to be elucidated, but it suggests that there is a connection between the potential transcriptional activity of a gene and the level of H3K27me3 required to keep it silenced.

In conclusion, we provide a CUT&Tag protocol applicable for small amount of tissues that works reliable as confirmed by comparing our data with previously published ChIP-seq data. The high signal-to-noise ratio typical for CUT&Tag allowed us to reduce the starting material substantially, enabling us to generate allele-specific TF binding profiles from early endosperm tissue, further the understanding of PHE1 transcriptional control and its relevance for seed development. These findings bridge transcription factor function with epigenetic regulation, offering new insights into how MADS-box proteins connect with imprinting mechanisms to control seed development.

## Methods

### Growth condition

The Col-0 *PHE1::PHE1-GFP* line was previously described by Batista et al. (2019). Seeds were sterilized by immersing them in 70% ethanol for 10 minutes, followed by 100% ethanol for 5 minutes. The ethanol was then removed, and the seeds were air-dried under sterile conditions. Sterilized seeds were sown on ½ strength Murashige and Skoog medium containing 1% sucrose and 0.68% agar. Stratification was performed in darkness at 4°C for two days. Seedlings were grown on plates for ten days under long-day conditions (16 hours of light at 21°C and 8 hours of darkness at 18°C with a light intensity of 150 µmol/s/m²) before being transferred to soil. Following transfer, plants were grown under long-day conditions with a light intensity of 150 µmol/s/m² and 70% relative humidity.

For experiments on homozygous plants siliques from self-fertilized Col-0 *PHE1::PHE1-GFP* plants were removed, leaving only open flowers corresponding to a time point of 1-2 days after pollination (DAP). The next day, siliques were harvested, weighed, flash-frozen in liquid nitrogen, and stored at -70°C. Material from approximately 2-3 plants were combined per replicate, each replicate consisted out of 250 mg of siliques.

For crossing experiments, five-week-old plants were emasculated and pollinated 48 hours later, as indicated. Siliques were harvested at 4 DAP, flash-frozen in liquid nitrogen, and stored at -70°C until nuclei isolation.

### Nuclei Isolation and CUT&Tag

Nuclei isolation was performed using approximately 250 mg of siliques per sample and replicate, utilizing the GentleMACS (Miltenyi Biotec) system as previously described (Moreno-Romero et al., 2017). A modified Honda buffer, in which MgCl_₂_ was replaced by 0.5 mM spermidine, was used to prevent carryover and potential activation of the transposase in subsequent steps. Following filtration and centrifugation for 6 minutes at 1600 g, nuclei were washed with 4 ml of CUT&Tag Wash150 buffer (150 mM NaCl, 20 mM HEPES-Na pH 7.5, 0.5 mM spermidine, 1x protease inhibitor) and centrifuged for 3 minutes at 600 g and 4°C. The supernatant was discarded, and the nuclei pellet was washed once with 2 ml of Wash150 buffer, followed by another centrifugation at 600 g for 3 minutes at 4°C. The resulting nuclei pellet was resuspended in 1000 µl of Wash150 buffer with 0.01% digitonin (Dig150 buffer) and centrifuged in a 2 ml tube before being resuspended in X*100 µl of antibody buffer (Dig150 buffer + 2 mM EDTA, pH 8) in 1.5 ml tubes. Since nuclei immobilization by ConA beads was unsuccessful in several attempts (data not shown), possibly due to the presence of starch or sugar, all steps requiring buffer exchange or washing were done by centrifugation for 3 min at 600 g and 4°C (600 g,3’, 4°C) if not indicated otherwise. For each 100 µl, 2 µl of primary antibody was added (anti-GFP ab290, Abcam; anti-H3, H9289-200UL, Sigma-Aldrich). Samples were gently mixed by flicking, collected by a quick centrifugation for 30 seconds at 600 g, and incubated overnight at 4°C on a nutator. Following incubation, the tubes were centrifuged (600 g,3’, 4°C), and the supernatant was discarded. Then, 100 µl of Dig150 buffer containing 1:100 secondary antibody (anti-Rabbit-IgG, ABIN101961, Antibodies-Online) was added, and the nuclei pellet was resuspended by pipetting.

Nuclei were incubated on a nutator for 30 min and subsequently centrifuged (600 g,3’, 4°C), the supernatant were removed and the pellet washed twice with each 300 ul Dig150 buffer. Afterwards, the pellet was eluted in 60 ul CUT&Tag Dig300 buffer (300 mM NaCl, 20 mM HEPES-Na pH 7.5, 0.5 mM spermidine, 1x protease inhibitor, 0.01% Digitonin) including 3 ul pAG-Tn5 enzyme (15-1017, EpiCypher). Nuclei were incubated on a nutator for 1h at room temperature, followed by another centrifugation and two rounds of washing with CUT&Tag 300 wash buffer. Hereafter, pellet was eluted in 100 ul Tagmentation buffer (300 mM NaCl, 20 mM HEPES-Na pH 7.5, 0.5 mM spermidine, 1x protease inhibitor, 0.01% Digitonin, 10 mM MgCl_2_) followed by one hour incubation at 37°C in a thermoblock in which tagmentation of target DNA fragments is happening. After tagmentation nuclei were collected by centrifugation the supernatant removed and the pellet washed with 200 ul TAPS buffer (10 mM TAPS, 2 mM EDTA pH 8) and centrifuged. Afterwards, supernatant was removed and 10 ul of SDS Release buffer (SDS, TAPS, EDTA) was added. Nuclei were vortexed for 10’’ and incubated for 1h at 58°C. Afterwards, 30 ul of SDS-quench buffer (0.67% Triton-X100 in H2O) was added, together with each 4 ul 10 mM i5 and i7 index primers. The solution was transferred to PCR tubes and amplified using Phusion Polymerase (F534L, Thermo Fisher) for 17 cycles. The resulting libraries were purified with 1.3x SPRIselect beads (B23318, Beckman) and eluted in 24 ul dH2O. Library integrity and concentration was determined using TapeStation D5000 ScreenTapes. Sequencing was performed on NextSeq1000 (Illumina) in 50 bp paired-end mode or in 150 bp paired-end mode for allele-specific experiments. For non-allele specific samples, it was aimed for 20 Mio paired-end reads, for allele specific samples 50 Mio reads were aimed for.

### Bioinformatic analysis

Raw fastq reads were trimmed and quality filtered using ‘TrimGalore’ (Version 0.6.10) (https://github.com/FelixKrueger/TrimGalore) using the parameters ‘-q 20 --stringency 3 -- length 32 –paired’. Trimmed reads were mapped to a the TAIR10 genome using ‘Bowtie2’ (Version 2.5.2)(Langmead and Salzberg, 2012) using the options ‘-I 10 -X 700 --no-unal -- no-mixed --no-discordant --phred33’ in the case of *PHE1::PHE1-GFP* samples and Col-0 isogenic wildtype samples. Alignment files were converted to bam files and quality filtered (-q 20) and only alignments kept that align to the nuclear chromosomes using ‘Samtools’ (Version 1.19.2)(Li et al., 2009). Filtered alignments were used for subsequent analysis.

Genome tracks where generated using deeptools ‘bamCoverage’ (Version 3.5.4.post1) (Ramírez et al., 2014) with the options -bs 50 --effectiveGenomeSize 119481543 -- normalizeUsing RPGC –skipNAs. Log2 normalization with of H3K27me3 to H3 was achieved using deeptools ‘bamCompare’ using the options --operation log2 -bs 10. Sample correlation matrices were obtained using either deeptools ‘multiBamSummary bins’ with the option -bs 5000 for genome wide analysis, or ‘multiBamSummary BED-file’ for target site specific correlation. Spearman correlation was obtained using deeptools ‘plotCorrelation’ with the options --corMethod spearman --whatToPlot scatterplot/heatmap --skipZeros –plotNumbers. Deeptools was run in a Python environment (Version 3.12.0)(Python Software, 2026). Quantification of signal in genomic intervals was achieved via bedtools map (Version 2.30) (Quinlan and Hall, 2010) using the -o mean operator.

Prior peak calling and to account for differences in sequencing depth between biological replicates, aligned CUT&Tag libraries were randomly downsampled to the sequencing depth of the replicate with the lowest number of properly paired alignments. The number of properly paired alignments was determined using SAMtools (samtools view -c -f 0x2). For each replicate, the required sampling fraction was calculated relative to the replicate with the lowest read count, and BAM files were randomly downsampled using samtools view -b -s with a fixed random seed of 42. Peaks were called using MACS2 callpeak (Version 2.2.9.1) (Zhang et al., 2008) with the parameters ‘--format BAMPE --gsize 1.119e8 -q 1e-3 in narrowPeak format for reproducibility analysis. Reproducibility peaks between biological replicates was assessed using the irreproducible discovery rate (IDR)(Li et al., 2011) framework (IDR v2.0.4.2). IDR analysis was performed for all pairwise combinations of biological replicates using the corresponding MACS2 narrowPeak files, with peaks ranked according to their p-value. Reproducible peaks were selected using a local IDR threshold of <0.05. Peaks passing this IDR reproducibility threshold were retained from the pairwise comparisons and combined to generate a final consensus set of high-confidence reproducible binding sites. Coordinates of the consensus peaks were used to annotate target regions with genic features using the package “ChIPseeker”(Yu et al., 2015) in R Version (4.3.3)(R Core Team, 2020).

Overlap with TE superfamilies from the Araport11 annotation (https://doi.org/10.6084/m9.figshare.30383395) as well as with ChIP-seq PHE1-GFP peak (Batista et al., 2019) was determined using Bedtools intersect (Version 2.30)(Quinlan and Hall, 2010) using the -u option.

PHE1 DNA-binding motifs were identified from 994 *PHE1::PHE1–GFP* CUT&Tag peak regions associated with genes using HOMER’s findMotifsGenome.pl (Version 5.1) (Heinz et al., 2010) function, using the default settings. To determine possible allele-specificity we performed CUT&Tag against PHE1-GFP from siliques at 4DAP in three independent replicates. Crossing were done using either Ler-0 wild type or *pistillata* maternal plants to generate the needed input of about 250 mg of siliques/replicate. All plants were pollinated with different individuals of the afore described Col-0 *PHE1::PHE1-GFP* line. CUT&Tag procedure was performed as described above, and subsequent sequencing of libraries was performed in 150 bp paired-end mode to increase the coverage of possible polymorphisms. Trimming of reads hereafter was done as described before. For allele specific analysis samples were mapped against a genome previously masked for SNPs between Col-0 (TAIR10) and Ler-0 using Bowtie2 as described above. The same subsequent quality-filtering steps were applied as described above. For visualization and to increase the sensitivity of detecting parentally biased binding events, allele-specific replicates were downsampled to match the fragment count of the replicate with the fewest fragments and subsequently merged. Allele discrimination was done with ‘SNPsplit’ (Version 0.6.0) using the –paired parameter (Krueger and Andrews, 2016). For subsequent analysis only reads specific to either one of the two genomes were used further, non-assigned of ambiguously-assigned reads were discarded. Reads in within the genomic coordinates of the afore defined 994 target regions associated with genes were counted using ‘featureCounts’ from the Subread package (Version 2.0.6) (Liao et al., 2014) with the options “-T 8 -p -O -s 0”. To avoid low coverage allele-specific reads we only considered those with >= 5 informative reads (reads that span a SNP) for each of the targets. Maternal ratio for those regions were calculated by Ler-reads/(Ler-reads + Col-reads). Cumulative CPM was calculated by the number of counts for a certain allele divided by counts from both allele i.e.(Col-reads/(Col-reads+Ler-reads)*1000000.

Publicly available mRNA-seq data was reanalyzed by downloading relevant samples from the corresponding archives. Raw reads were trimmed using Trim Galore (Version 0.6.10) (Krueger, 2015) with the options ‘--fastqc --clip_R1 5 --three_prime_clip_R1 20’. Trimmed reads were mapped against the TAIR10 genome or for allele-specific analysis against a Col-Ler masked version of the TAIR10 genome using Hisat2 (Version 2.2.1)(Kim et al., 2019) with the ‘–dta’ parameter set. Reads were counted using ‘featureCounts’ (Version 2.0.6) (Liao et al., 2014) with the options ‘-T 10 -t exon -g gene_id’. Downstream expression analysis was performed with DESeq2 (Love et al., 2014). Allele-specific analysis was done as described above for CUT&Tag derived fragments using >5 informative reads/gene.

Statistical analysis was conducted using R (Version 4.3.3. and version 4.4.2) (R Core Team, 2020) as indicate per plot and plots were generated using R base or the ‘ggplot2’ package (Wickham, 2009). Final arrangement of Figures was done with Affinity Designer 2 (Version 2.1.1; Serif Europe).

## Data availability

CUT&Tag data generated for this study was uploaded to Gene Expression Omnibus (https://www.ncbi.nlm.nih.gov/geo/) under the accession number GSE355283. Allele-specific transcriptome data from 4DAP was taken from (Moreno-Romero et al., 2019). Allele-specific expression data in *fie* and wild type background and wild type from (Hsieh et al., 2011) accession number GSE24644. Allele-specific H3K27me3 and DNA methylation data was reanalyzed from (Del Toro-De Leon et al., 2024) available under GEO accession number GSE260822.

## Acknowledgements

This work was financially supported by the Max Planck Society. Writing and coding for this manuscript was assisted provided by an AI-powered chat service, hosted by the Göttingen University’s Gesellschaft für wissenschaftliche Datenverarbeitung (GWDG). ChatGPT (OpenAI, GPT-6) was used to assist with manuscript language editing and the development and debugging of computational analysis scripts. The authors independently verified the resulting analyses and take full responsibility for the content of the manuscript.

---

**Supplementary Figure S1:**
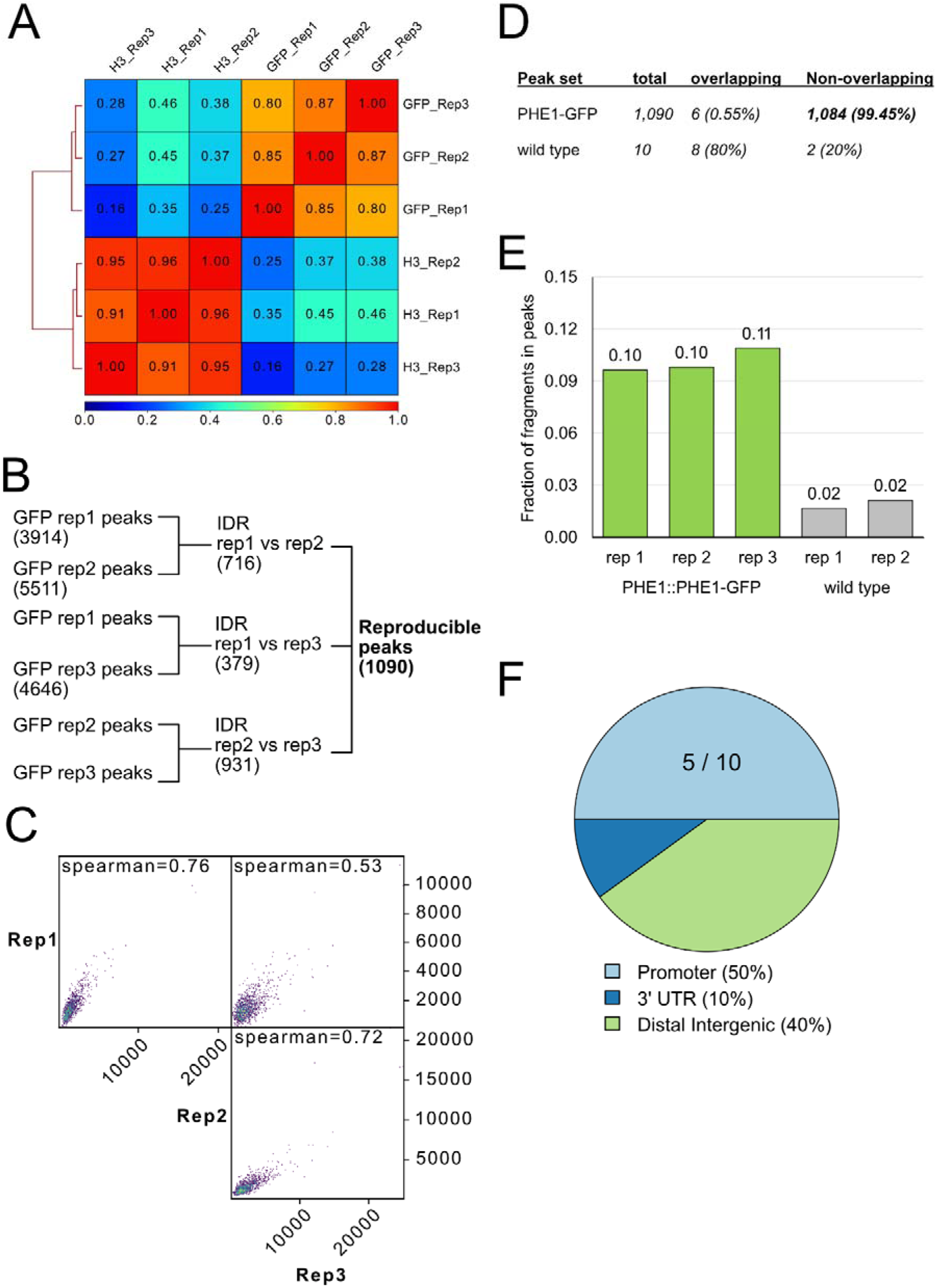
A) Heatmap of Spearman correlation of from CUT&Tag samples used for ChIP validation. Anti-H3 samples are labeled with “H3” anti-GFP samples with “GFP”, replicates are indicated by “Rep”. B) Schematic representation of irreproducible discovery rate (IDR) workflow. For this mapped GFP samples were down-sampled to the read number of the smallest replicate and subsequently peaks were called for each replicate separately using MACS2 with a relaxed threshold (p < 0.001). All three pairwise replicate comparisons were subjected to IDR analysis, and peaks with a global IDR ≤0.05 were retained. Overlapping reproducible regions were subsequently merged to obtain the final consensus peak set. Numbers in brackets show the number of peaks meeting the described thresholds per set. C) Scatterplot of Spearman correlation within the genomic coordinates of reproducible peaks (n= 1090). D) Number of peaks detected in Col-0 *PHE1::PHE1-GFP* samples and transgene-free wild type samples, as well as the reciprocal overlap calculated by bedtools intersect. Eight of the ten background peaks overlapped six distinct PHE1 peaks, as multiple background peaks could overlap the same PHE1 region. E) Fraction of fragments in peaks (FFiPs) of the three replicates form *PHE1::PHE1-GFP* containing samples and the two transgene-free Col-0 replicates. FFiP was calculated over al 1090 peaks regions as defined above. F) Feature annotation of the ten peaks from transgene-free samples done with ChIPseeker. Number of peaks in promoter are plotted.

**Supplementary Figure S2:**
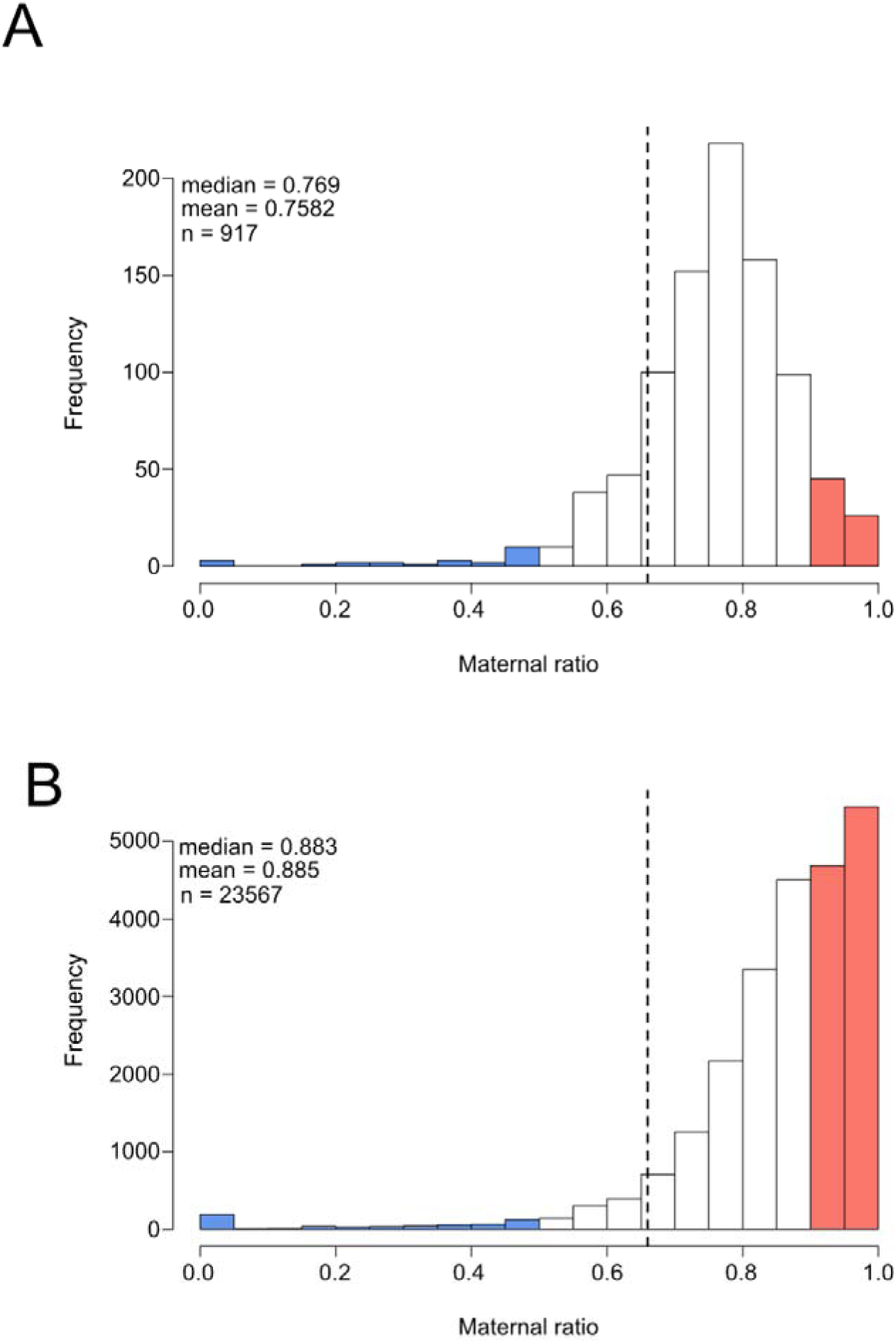
Maternal ratio distribution of PHE1-GFP binding in defined promoter windows of -1500bp to +500 bp relative to genes. A) Histogram of maternal ratios promoters of PHE1 target genes. B) Histogram of maternal ratios in all Araport11 annotated promoters in the afore defined windows. Blue represents all promoters with a bias for paternal PHE1 binding (< 0.5), red maternal PHE1 binding (>0.9). Dashed lines represent the theoretical allele distribution of 0.66 in the endosperm. Shown are only promoters with >= 5 informative reads, n, mean and median as indicated.

**Supplementary Figure S3:**
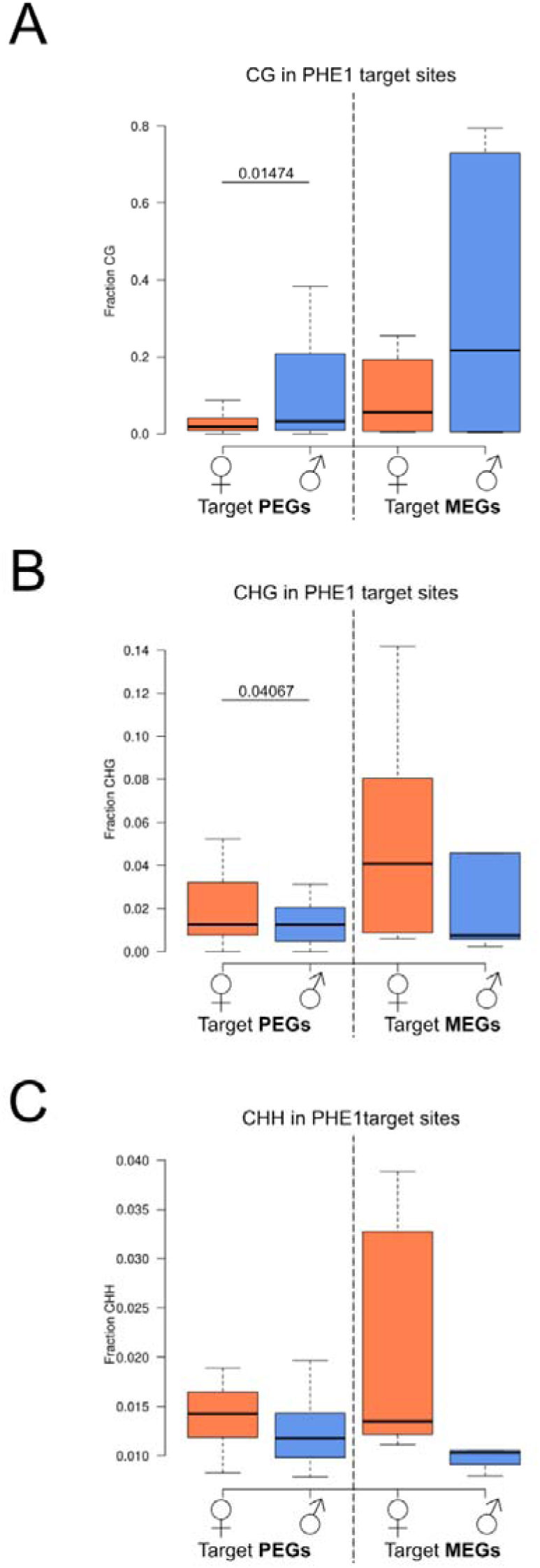
Average DNA methylation per context in PHE1 target sites. DNA methylation is shown in CG (A), CHG (B) and CHH (C) whereas H represents any nucleotide other than G. Analysis was performed over all PHE1 targeted PEGs (n=20) or MEGs (n=6) respectively. Orange represents the maternal allele (♀), blue the paternal alleles (♂). Statistical analysis between same parental alleles between MEGs and PEGs was performed using unpaired Wilcoxon rank-sum test, whereas different alleles from MEGs, or PEGs were tested via paired Wilcoxon signed-rank test. Significant p-values are plotted above. DNA methylation data from combined Ler × Col replicates from Del Toro-De León et al. 2024.

**Supplementary Figure S4:**
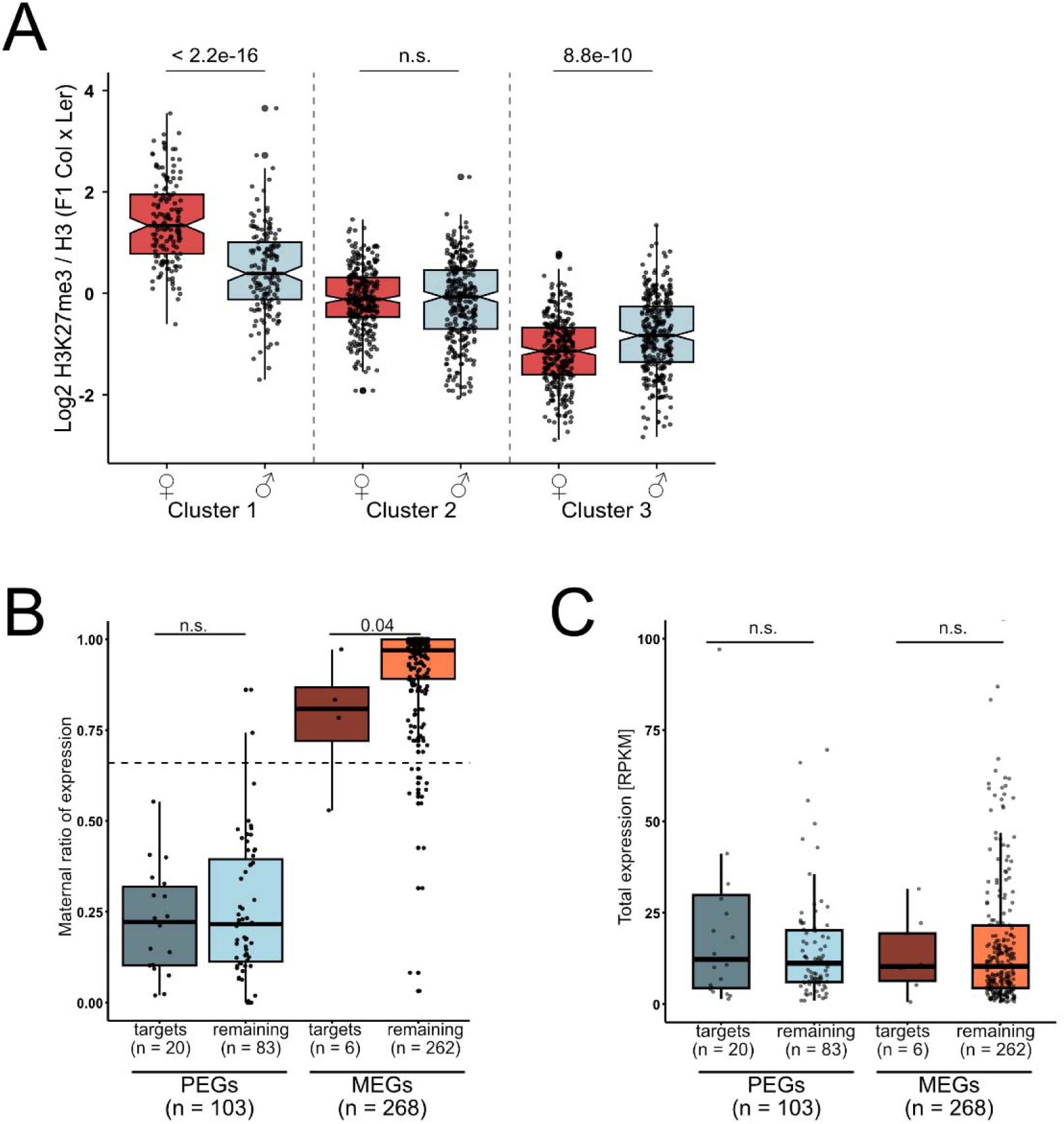
A) Parental specific H3K27me3 in clusters as defined before, reciprocal to Fig 3E. H3K27me3 data are from Del Toro-De León et al. 2024. Expression data is 4 DAP INTACT purified endosperm from F1 Ler × Col crosses from Moreno-Romero et al. 2019. B) Maternal ratio of expression. C) Total expression of target genes in RPKM. Statistical significance was determined using unpaired Wilcoxon rank-sum test.

## References

1. Dilkes, B. P., Spielman, M., Weizbauer, R., Watson, B., Burkart-Waco, D., Scott, R. J. & Comai, L. 2008. The Maternally Expressed WRKY Transcription Factor Controls Lethality in Interploidy Crosses of Arabidopsis. Plos Biology, 6, 2707–2720.

2. Scott, R. J., Spielman, M., Bailey, J. & Dickinson, H. G. 1998. Parent-of-origin effects on seed development in. Development, 125, 3329–3341.

3. Gehring, M., Missirian, V. & Henikoff, S. 2011. Genomic Analysis of Parent-of-Origin Allelic Expression in Seeds. Plos One, 6.

4. Hsieh, T. F., Shin, J. Y., Uzawa, R., Silva, P., Cohen, S., Bauer, M. J., Hashimoto, M., Kirkbride, R. C., Harada, J. J., Zilberman, D. & Fischer, R. L. 2011. Regulation of imprinted gene expression in Arabidopsis endosperm. Proceedings of the National Academy of Sciences of the United States of America, 108, 1755–1762.

5. Batista, R. A. & Köhler, C. 2020. Genomic imprinting in plants-revisiting existing models. Genes Dev, 34, 24–36.

6. Gehring, M., Bubb, K. L. & Henikoff, S. 2009. Extensive Demethylation of Repetitive Elements During Seed Development Underlies Gene Imprinting. Science, 324, 1447–1451.

7. Pignatta, D., Erdmann, R. M., Scheer, E., Picard, C. L., Bell, G. W. & Gehring, M. 2014. Natural epigenetic polymorphisms lead to intraspecific variation in Arabidopsis gene imprinting. Elife, 3.

8. Moreno-Romero, J., Jiang, H., Santos-Gonzalez, J. & Kohler, C. 2016. Parental epigenetic asymmetry of PRC2-mediated histone modifications in the Arabidopsis endosperm. EMBO J, 35, 1298–311.

9. Mozgova, I., Köhler, C. & Hennig, L. 2015. Keeping the gate closed: functions of the polycomb repressive complex PRC2 in development. Plant Journal, 83, 121–132.

10. Zhang, M., Xie, S. J., Dong, X. M., Zhao, X., Zeng, B., Chen, J., Li, H., Yang, W. L., Zhao, H. N., Wang, G. K., Chen, Z. L., Sun, S. L., Hauck, A., Jin, W. W. & Lai, J. S. 2014. Genome-wide high resolution parental-specific DNA and histone methylation maps uncover patterns of imprinting regulation in maize. Genome Research, 24, 167–176.

11. Kaya-Okur, H. S., Wu, S. J., Codomo, C. A., Pledgers, E. S., Bryson, T. D., Henikoff, J. G., Ahmad, K. & Henikoff, S. 2019. CUT&Tag for efficient epigenomic profiling of small samples and single cells. Nature Communications, 10.

12. Weinhofer, I., Hehenberger, E., Roszak, P., Hennig, L. & Köhler, C. 2010. H3K27me3 Profiling of the Endosperm Implies Exclusion of Polycomb Group Protein Targeting by DNA Methylation. Plos Genetics, 6.

13. Batista, R. A., Moreno-Romero, J., Qiu, Y., Van Boven, J., Santos-Gonzalez, J., Figueiredo, D. D. & Köhler, C. 2019. The MADS-box transcription factor PHERES1 controls imprinting in the endosperm by binding to domesticated transposons. Elife, 8.

14. Abbasova, L., Urbanaviciute, P., Hu, D., Ismail, J. N., Schilder, B. M., Nott, A., Skene, N. G. & Marzi, S. J. 2025. CUT&Tag recovers up to half of ENCODE ChIP-seq histone acetylation peaks. Nat Commun, 16, 2993.

15. Yu, C. P., Lin, J. J. & Li, W. H. 2016. Positional distribution of transcription factor binding sites in Arabidopsis thaliana. Sci Rep, 6, 25164.

16. Heyndrickx, K. S., Van De Velde, J., Wang, C., Weigel, D. & Vandepoele, K. 2014. A functional and evolutionary perspective on transcription factor binding in Arabidopsis thaliana. Plant Cell, 26, 3894–910.

17. O’malley, R. C., Huang, S. C., Song, L., Lewsey, M. G., Bartlett, A., Nery, J. R., Galli, M., Gallavotti, A. & Ecker, J. R. 2016. Cistrome and Epicistrome Features Shape the Regulatory DNA Landscape. Cell, 166, 1598.

18. Del Toro-De Leon, G., Van Boven, J., Santos-Gonzalez, J., Jiao, W. B., Peng, H., Schneeberger, K. & Kohler, C. 2024. Epigenetic and transcriptional consequences in the endosperm of chemically induced transposon mobilization in Arabidopsis. Nucleic Acids Res.

19. Moreno-Romero, J., Del Toror-De León, G., Yadav, V. K., Santos-González, J. & Köhler, C. 2019. Epigenetic signatures associated with imprinted paternally expressed genes in the Arabidopsis endosperm. Genome Biology, 20.

20. Grossniklaus, U., Vielle-Calzada, J. P., Hoeppner, M. A. & Gagliano, W. B. 1998. Maternal control of embryogenesis by Medea, a Polycomb Group Gene in Arabidopsis. Science, 280, 446–450.

21. Köhler, C., Hennig, L., Bouveret, R., Gheyselinck, J., Grossniklaus, U. & Gruissem, W. 2003. Arabidopsis MSI1 is a component of the MEA/FIE Polycomb group complex and required for seed development. EMBO J, 22, 4804–14.

22. Bartosovic, M., Kabbe, M. & Castelo-Branco, G. 2021. Single-cell CUT&Tag profiles histone modifications and transcription factors in complex tissues. Nature Biotechnology, 39, 825– 835.

23. Zheng, X. Y. & Gehring, M. 2019. Low-input chromatin profiling in Arabidopsis endosperm using CUT&RUN. Plant Reproduction, 32, 63–75.

24. Tao, X. Y., Feng, S. L., Zhao, T. & Guan, X. Y. 2020. Efficient chromatin profiling of H3K4me3 modification in cotton using CUT&Tag. Plant Methods, 16.

25. Kaya-Okur, H. S., Janssens, D. H., Henikoff, J. G., Ahmad, K. & Henikoff, S. 2020. Efficient low-cost chromatin profiling with CUT&Tag. Nat Protoc, 15, 3264–3283.

26. Gehring, M. 2013. Genomic imprinting: insights from plants. Annu Rev Genet, 47, 187–208.

27. Rodrigues, J. A. & Zilberman, D. 2015. Evolution and function of genomic imprinting in plants. Genes Dev, 29, 2517–31.

28. Köhler, C., Page, D. R., Gagliardini, V. & Grossniklaus, U. 2005. The Arabidopsis thaliana MEDEA Polycomb group protein controls expression of PHERES1 by parental imprinting. Nat Genet, 37, 28–30.

29. Bemer, M., Heijmans, K., Airoldi, C., Davies, B. & Angenent, G. C. 2010. An atlas of type I MADS box gene expression during female gametophyte and seed development in Arabidopsis. Plant Physiol, 154, 287–300.

30. De Folter, S., Immink, R. G., Kieffer, M., Parenicova, L., Henz, S. R., Weigel, D., Busscher, M., Kooiker, M., Colombo, L., Kater, M. M., Davies, B. & Angenent, G. C. 2005. Comprehensive interaction map of the Arabidopsis MADS Box transcription factors. Plant Cell, 17, 1424–33.

31. Jadhav, U., Manieri, E., Nalapareddy, K., Madha, S., Chakrabarti, S., Wucherpfennig, K., Barefoot, M. & Shivdasani, R. A. 2020. Replicational Dilution of H3K27me3 in Mammalian Cells and the Role of Poised Promoters. Mol Cell, 78, 141–151 e5.

32. Moreno-Romero, J., Santos-Gonzalez, J., Hennig, L. & Kohler, C. 2017. Applying the INTACT method to purify endosperm nuclei and to generate parental-specific epigenome profiles. Nat Protoc, 12, 238–254.

33. Langmead, B. & Salzberg, S. L. 2012. Fast gapped-read alignment with Bowtie 2. Nat Methods, 9, 357–9.

34. Li, H., Handsaker, B., Wysoker, A., Fennell, T., Ruan, J., Homer, N., Marth, G., Abecasis, G., Durbin, R. & Proc, G. P. D. 2009. The Sequence Alignment/Map format and SAMtools. Bioinformatics, 25, 2078–2079.

35. Ramírez, F., Dündar, F., Diehl, S., Grüning, B. A. & Manke, T. 2014. deepTools: a flexible platform for exploring deep-sequencing data. Nucleic Acids Research, 42, W187–W191.

36. PYTHON Software, F. 2026. Python Programming Language. Python Software Foundation.

37. Quinlan, A. R. & Hall, I. M. 2010. BEDTools: a flexible suite of utilities for comparing genomic features. Bioinformatics, 26, 841–842.

38. Zhang, Y., Liu, T., Meyer, C. A., Eeckhoute, J., Johnson, D. S., Bernstein, B. E., Nusbaum, C., Myers, R. M., Brown, M., Li, W. & Liu, X. S. 2008. Model-based analysis of ChIP-Seq (MACS). Genome Biol, 9, R137.

39. Li, Q., Brown, J. B., Huang, H. & Bickel, P. J. 2011. Measuring reproducibility of high-throughput experiments.

40. Yu, G. C., Wang, L. G. & He, Q. Y. 2015. ChIPseeker: an R/Bioconductor package for ChIP peak annotation, comparison and visualization. Bioinformatics, 31, 2382–2383.

41. R CORE Team, R. 2020. R: A language and environment for statistical computing.

42. Citeseer Heinz, S., Benner, C., Spann, N., Bertolino, E., Lin, Y. C., Laslo, P., Cheng, J. X., Murre, C., Singh, H. & Glass, C. K. 2010. Simple combinations of lineage-determining transcription factors prime cis-regulatory elements required for macrophage and B cell identities. Mol Cell, 38, 576–89.

43. Krueger, F. & Andrews, S. R. 2016. SNPsplit: Allele-specific splitting of alignments between genomes with known SNP genotypes. F1000Res, 5, 1479.

44. Liao, Y., Smyth, G. K. & Shi, W. 2014. featureCounts: an efficient general purpose program for assigning sequence reads to genomic features. Bioinformatics, 30, 923–930.

45. Krueger, F. 2015. Trim Galore!: A wrapper around Cutadapt and FastQC to consistently apply adapter and quality trimming to FastQ files, with extra functionality for RRBS data. Babraham Institute.

46. Kim, D., Paggi, J. M., Park, C., Bennett, C. & Salzberg, S. L. 2019. Graph-based genome alignment and genotyping with HISAT2 and HISAT-genotype. Nat Biotechnol, 37, 907–915.

47. Love, M. I., Huber, W. & Anders, S. 2014. Moderated estimation of fold change and dispersion for RNA-seq data with DESeq2. Genome Biol, 15, 550.

48. Wickham, H. 2009. ggplot2: Elegant Graphics for Data Analysis. Ggplot2: Elegant Graphics for Data Analysis, 1–212.

